# Perceiving Hesitation requires both the Motor and Mentalizing systems

**DOI:** 10.1101/454298

**Authors:** Marc Thioux, Judith Suttrup, Christian Keysers

## Abstract

The mentalizing network and the putative mirror neuron system (pMNS) are two important large scale brain networks for social cognition, with very little overlap between them. Evidence suggests however that the two networks can collaborate for understanding the state of mind of others on the basis of their body movements. Here using fMRI we find that when participants view hand actions to detect hesitations they activate both their mentalizing and pMNS networks and information is exchanged across these networks. In a follow-up experiment using repetitive TMS we find that disturbing activity in either network slows hesitation attribution. In contrast, watching the same actions to determine the size of the object being grasped only triggered activity in the pMNS network, and rTMS over the mentalizing network then no longer slowed reactions. When we see others’ actions, we recruit our own motor system. Our results suggest that for detecting a simple mental state like hesitation, this pre-processed motor information becomes a necessary input into a mentalizing network that is essential for associating deviations from a predicted motor program with a specific mental state.

## INTRODUCTION

Understanding non-verbal behaviours is essential for social interactions as it can provide information about the state of mind of others, notably their hidden intentions, beliefs, and emotional states. Sometimes, these states of mind are betrayed by very subtle kinematic cues like the raise of an eye-brow or a pause during a reach-to-grasp action.

A great number of studies have investigated the neural basis of the ability to reflect about other people’s mental states (Mar 2011; Schurz et al. 2014). One of the most investigated mental states are false-beliefs. The ability to understanding that someone else has a false belief is strong evidence for the presence of the capacity to read other’s mind as it requires dissociating one’s own perspective on reality from the one of the other. One way to study the neural basis of the ability to understand mental states has therefore been to present participants with stories involving a protagonist who has a false belief and compare the brain activity with that while presenting similar stories that do not involve mental state attribution but require for instance to recognize that a map does not represent the current reality (Saxe and Wexler 2005). Understanding false-belief stories is associated with increased activity in a mentalizing network including the medial prefrontal cortices (mPFC), temporoparietal junctions (TPJ), temporal poles, superior temporal sulci (STS), precuneus (PreCu) and anterior inferior frontal gyri (aIFG) (Blue in Figure 1). The same network has been shown to be activated by a variety of stimuli like cartoons, photographs of the eye, or movies of geometrical shapes behaving like humans, and for a variety of mental states like beliefs, intentions and desires (Mar 2011; Schurz et al. 2014).

**Figure 1:**
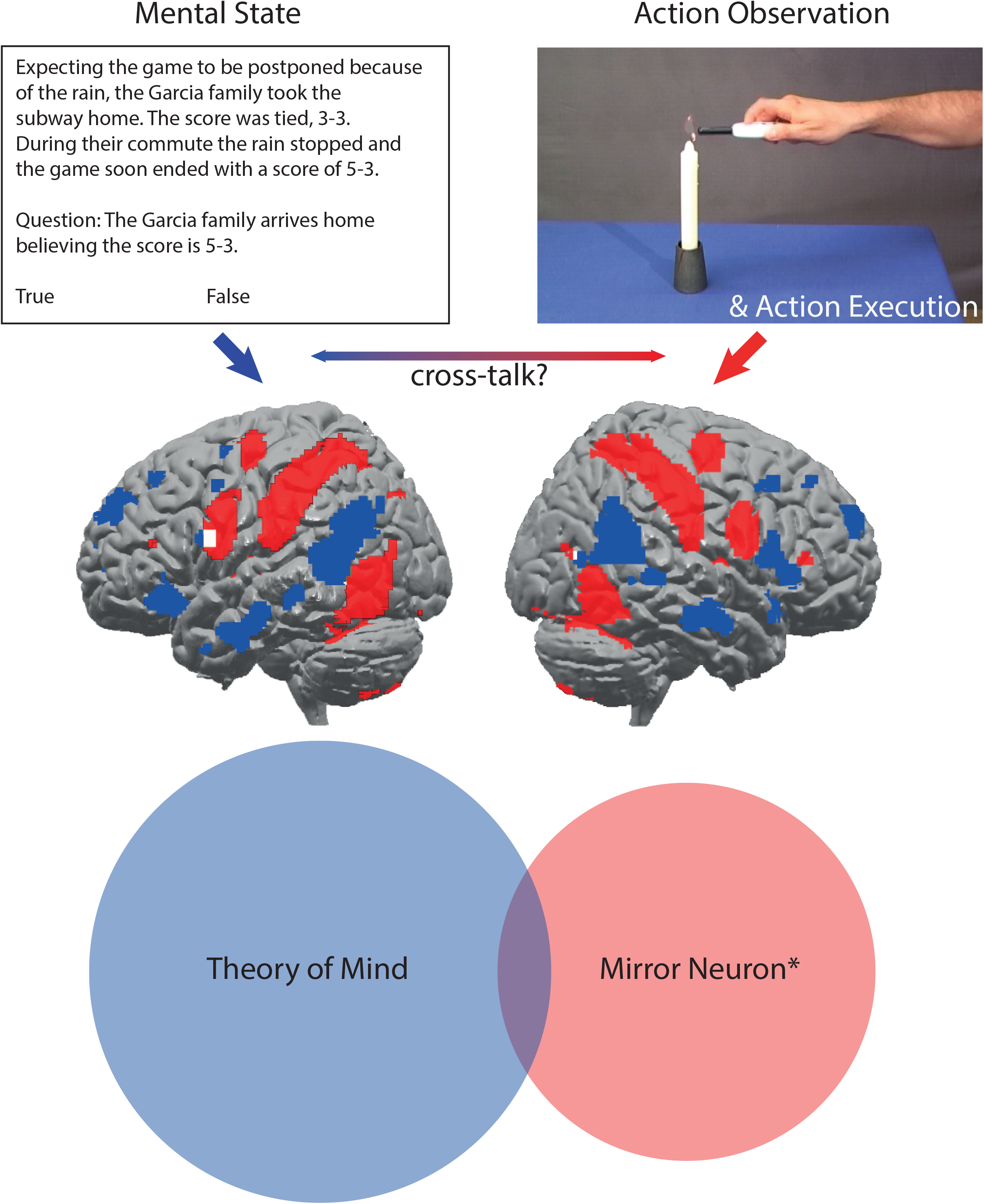
Top: Two fields of research have investigated networks involved in processing the inner states of others: one investigates how we attribute mental states to others (left, e.g using short stories from Saxe & Wexler (2005)) and identified the blue network (Mar (2011)); the other how humans and monkeys process observed actions (e.g. video from Arnstein et al (Arnstein et al. 2011)) and map voxels/cells involved both in the observation of actions and the execution of similar actions (red network in the render, (Gazzola and Keysers 2009)). Intriguingly very little overlap between these networks is observed (white). Bottom: A web of science search confirms the limited overlap between neuroscience literature on Theory of Mind (blue) and Mirror Neurons (red) using search terms “Theory of Mind” and “Mirror Neuron*” as topics, respectively and jointly (overlap), as illustrated by this Venn diagram where surface is proportional number of papers in the field of Neuroscience.

Seeing body movements such as goal-directed hand actions is associated with increased activity in a set of brain regions that show conspicuously little overlap with the mentalizing network (red network in Figure 1). These include visual areas specialized in processing body shape and biological motion (V5 and posterior STS), as well as several brain regions also recruited when performing similar actions, including parietal region PF, the ventral premotor cortex (vPM), the primary somatosensory cortex (SI), the dorsal premotor cortex (dPM) (Gazzola and Keysers 2009; Keysers and Gazzola 2009; Caspers et al. 2010; Keysers et al. 2010; Molenberghs et al. 2012). These later brain regions will be called the putative mirror neuron system (pMNS) because of their homologies with the monkey mirror neuron system, where the qualifier ‘putative’ acknowledges that a region can show activations during action observation and execution without containing mirror neurons (Gazzola and Keysers 2009). Lesion and brain stimulation studies suggest that the re-enactment of others action in the pMNS supports certain aspects of action comprehension (Avenanti et al. 2013; Urgesi et al. 2014; Keysers et al. 2018).

As demonstrated by meta-analyses, the mentalizing network is generally not activated in studies of action observation, and actually the two networks have been found to be anti-correlated at rest (Uddin et al. 2009), and during the inhibition of imitation (Brass et al. 2009). Nonetheless, several studies have reported increases of activity in the mentalizing network while subjects watched actions performed by a protagonist. Results of these studies suggest that increases of activity or connectivity in the mentalizing system is found when participants watch interactive or communicative gestures (Schippers et al. 2009; Centelles et al. 2011; Becchio et al. 2012; Mainieri et al. 2013; Ciaramidaro et al. 2014; Sperduti et al. 2014) when watching irrational actions in relation to the context (Brass et al. 2007; Jastorff et al. 2011; but see Liew et al. 2011; Ampe et al. 2014) or when simply paying attention to *why* an action is performed (Brunet et al. 2000; Grèzes et al. 2004; de Lange et al. 2008; Spunt et al. 2011; Chambon et al. 2017)

This tension between studies emphasizing the independence of these two networks and those reporting co-activation raises the question of what determines whether observed actions recruit the theory of mind network. Several attempts have been made to describe the different levels of representations involved in actions, and these distinctions might be relevant here. Searle distinguishes intention-in-action and prior intention: “suppose I have a prior intention to drive to my office. As I am carrying out this prior intention I might perform a series of subsidiary actions for which I need not have formed a prior intention: opening the door, starting the engine, depressing the clutch, etc. When I formed my intention to drive to the office I might not have given these subsidiary acts a thought. Yet such actions are intentional. For such cases I have an intention, but no prior intentions. All intentional actions have intentions-in-action but not all intentional actions have prior intentions. “ (p52,Searle 1980). Others have distinguished three levels of motor representations.Grafton and Hamilton distinguish kinematics, object-goals and outcomes (Grafton and Hamilton 2007), and we have distinguished the how, what and why of actions: “The professor leans back into his chair and reaches with his left hand for one of the books on the table. Your perception of this last action has at least three levels. At the lowest, most detailed level, you could perceive *how* he performed this action: with his left hand, using a whole hand prehension. At the intermediate level, you perceive *what* he is doing: grasping a book. At the highest level you might perceive *why* he is doing it: to signal that your conversation is over.” (Thioux et al. 2008). Could it be that tasks that encourage participants to think about more abstract representations (prior intentions and why) and use stimuli that omit the kinematics of a bodily action trigger mentalizing brain activity while tasks that emphasize bodily kinematics and do not encourage reflecting about more abstract levels only activate the pMNS (Gobbini et al. 2007; Keysers and Gazzola 2007; Spunt and Lieberman 2012). A way to test this possibility is to show participant movies depicting bodily kinematics under two instructions: one focusing on less abstract and one focusing on more abstract mental states. We would then expect the two systems to be coactivated in the latter but not the former condition.

But what exactly should that more abstract mental state be? If this mental state is too abstract and detached from the kinematics of the action, reflecting about the mental state may recruit the mentalizing system and make the pMNS irrelevant (Kilner et al. 2007; Kilner and Frith 2008). Here we therefore explore a mental state closer to observable action kinematics: confidence. Participants can judge how confident someone is based on the kinematics of simple observed goal-directed actions (Patel et al. 2012), providing a potential for a link with the pMNS. At the same time, hesitations are more abstract than the intentions-in-action that are often found to recruit the pMNS alone (e.g. which of two objects is this hand aiming to grasp?).

We therefore showed participants an actor reaching into a box to grasp one of two balls that are occluded by the box (Figure 2). One of the balls had a diameter of 10 cm, the other of 3.5 cm. In some stimuli, we instructed actors to grasp the small ball or the large ball, a very simple instruction that lead to confident grasping. In other stimuli, we instructed actors to grasp the more or less saturated of two balls with similar colour-saturation. This resulted in genuine hesitant grasp – that is to say, that actors did not aim to hesitate, but aimed to grasp the more or less saturated ball, and hesitations emerged from a difficulty to translate this prior intention into a specific intention-in-action (to grasp the right or left ball). Stimuli were then selected based on the fact that a majority of independent raters could recognize the size of the ball being grasped and the presence of hesitations. We then asked participants to perform two different tasks on these stimuli while measuring their brain activity with fMRI: either to judge whether the hand is grasping the large or small ball (Target task) or whether the actor is confident or hesitant (Hesitation task). We then found according to our intuition that (i) the Target task recruited the pMNS but not the mentalizing system, (ii) the Hesitation task recruited both the pMNS and mentalizing network. This is in line with a small number of fMRI studies showing co-activation of the two system (Grèzes et al. 2004; Schippers et al. 2009; Centelles et al. 2011; Spunt et al. 2011; Becchio et al. 2012; Mainieri et al. 2013; Ciaramidaro et al. 2014; Kuhlen et al. 2015). Functional connectivity analyses revealed significant functional connectivity across these networks, with information exchange between the mentalizing network and visual regions mediated (at least in part) by the pMNS. However, the most critical test of the notion that the pMNS and mentalizing systems can work together to inform social cognition would be to show that interfering with both systems interferes with the detection of Hesitations, for instance by slowing reaction times. To date, such causal evidence is missing. We therefore later performed a follow-up repetitive TMS study with new participants and show (iv) that interfering with activity in the pMNS slowed reaction times in both tasks (Target and Hesitation), while interfering with activity in the mentalizing network only slowed reaction times in the hesitation task.

**Figure 2:**
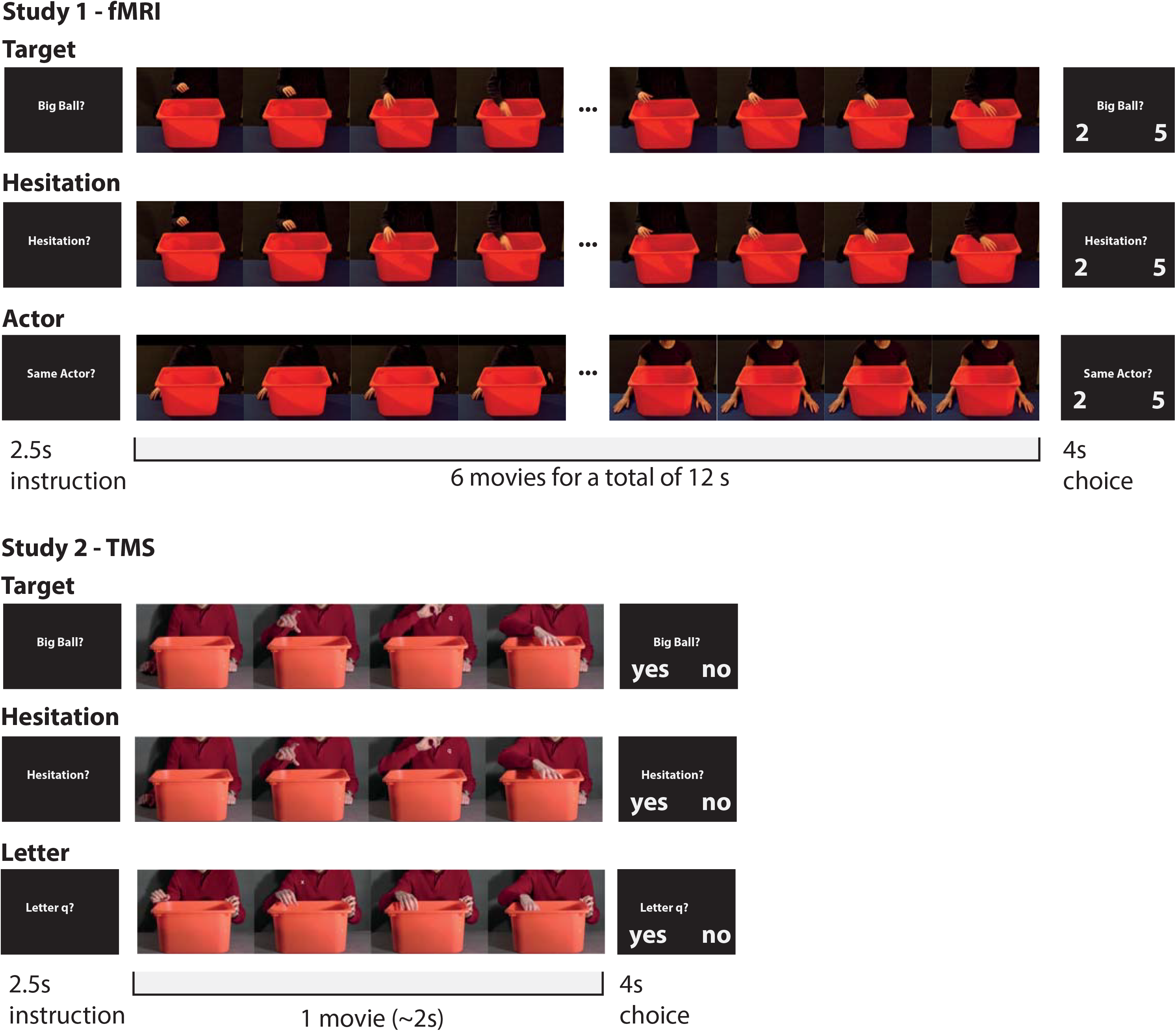
Study Design. In Study 1, participants’ brain activity was measured while they saw a series of 6 movies depicting an actor reach into a box, and needed to determine how often the actor grasped the big ball (Target condition), or how often the actor was hesitant (Hesitation condition), based on the same movies. In the control condition, the participants needed to determine how often they witnessed the same actor in the 6 still frames shown for the same duration as the movies in the other conditions (Actor condition). In Study two, participants were under the effect of rTMS over the rTPJ, rPF or vertex, while they had to determine whether the actor grasped the big ball (Target condition) or hesitated (Hesitation condition). In the control condition, participants had to indicate whether a small q or x appeared in the movie. In all cases, an instruction screen indicated participants what task they need to perform before the stimuli are presented to trigger the deliberate deployment of neural resources to that specific task.

In both studies, we needed control conditions. In the fMRI study (Study 1), we were keen to identify the basic network processing action kinematics, and thus needed a task similar to our experimental tasks but without kinematic information. We therefore presented still frames from our original stimuli to remove kinematic information and asked participants to detect changes in identity to nevertheless tax executive functions. In the TMS study, our main dependent variable was reaction time, and we thus needed a task without action processing that could control for unspecific changes in the motor system across sessions. We therefore asked participants to report a letter that was briefly superimposed on the original stimuli.

## METHODS

### General comments

The study is composed of two studies, with the TMS experiment (Study 2) performed over a year after the fMRI experiment (Study 1) to explore the causal contribution to mental state attribution of the systems found to be correlated with the task in the first study. The two studies were thus performed on different participants, as the participants of Study 1 could no longer be called back into the lab. Stimuli were also re-recorded for Study 2 to satisfy the specific requirements of a TMS reaction time study. In Study 1, to inform reverse inference, we additionally performed two localizer tasks to map brain regions involved in grasping and mentalizing. In Study 2, to neuro-navigate the TMS stimulation we also performed fMRI experiments with each participant to localize the mentalizing and pMNS system prior to the TMS experiment. Table 1 summarizes the number of participants involved in each experiment.

**Table 1:** Summary of experiments and participants involved in the two studies. For each part of the experiment we indicade, from left to right how many participants were included in the final analysis presented in the results section, their age (±std), how many participants had met the inclusion criteria (e.g. MRI compatibility for Study 1, and Passive Motor Threshold ≤72% of our machine output for Study 2), and how many were excluded from the final analysis. If the sub-experiments of each study involved the same participants, the cells are merged. The 14 participants of the functional connectivity study were all part of the 23 involved in the other parts of Study 1 but none of the 14 involved in Study 2 had participated in Study 1. Reasons for exclusion included (*) two from the 23 Target/Hesitation participants were excluded because they were not Dutch speaking and one because the task failed to run in the stimulus presentation due to technical issues. (**) 9 of the original 23 were excluded because not all 7 regions of interest could be identified in their single subject fMRI data, see methods; (***) one asked to discontinue the experiment after the first rTMS session, two had excessive head motion during the rTMS stimulation, one was not able to perform the behavioral task above chance level. (+) Originally, 35 participants responded to the invitation to participate in the study, but 17 failed to meet the inclusion criteria: one because of general TMS safety criteria, and 16 because we failed to trigger a motor response using 65% of the TMS system’s maximum power (using higher power would have led to overheating during the rTMS procedure and thus unreliable stimulation).

| Participants | in final analysis | age | Met Inclusion Criteria | excluded |
| --- | --- | --- | --- | --- |
| Study 1 |  |  |  |  |
| Target/Hesitation fMRI | 23 | 22 ± 3 | 23 | 0 |
| Action Execution fMRI |  |  |  |  |
| False-Belief fMRI | 20 |  |  | 3* |
| Functional Connectivity | 14 |  | 23 | 9** |
| Study 2 |  |  |  |  |
| False-Belief fMRI | 14 | 22 ± 2 | 18<sup>+</sup> | 4*** |
| Action Observation fMRI |  |  |  |  |
| Action Execution fMRI |  |  |  |  |
| Target/Hesitation TMS |  |  |  |  |

## STUDY-1: fMRI

### Participants

Twenty-three young adults (10 females and 13 males between 19 and 30 years of age; mean 22 ± 2.9) participated in the study. All were right handed according to the Edinburgh handedness inventory (mean handedness score 87.6% ± 12.8, min = 60, max = 100), and had normal or corrected-to-normal vision. None had a history of neurological or psychiatric disorder. Procedures were approved by the Institutional Review Board (medical ethical committee), and volunteers gave written informed consent to participate.

### MRI-scanning

The study comprised three experiments, in the same order for all participants: (1) the main Target/Hesitation experiment, (2) Action execution localizer, (3) False-beliefs understanding localizer. The study was carried out on a Philips 3T Intera MR-scanner equipped with a 32-channel head coil. Functional images were acquired parallel to the bi-commissural plane using a T2*-weighted single-shot echo-planar sequence with TR = 2.0 s, TE = 28 ms, and 39 axial slices 3.5 mm thick (with no gap) covering the whole brain (flip angle 70 degrees, FOV 224 x 136.5 x 224 mm, Reconstructed Matrix size 64 x 64, isometric 3.5 mm voxels). A T1-weighted anatomical image consisting of 160 slices 1mm-thick was acquired at the beginning of the study for normalization purpose (TR 7.6 ms, TE 3.5 ms, flip angle 8 degrees, FOV 224 x 160 x 256 mm, reconstructed matrix size 256 x 256, isometric 1 mm voxels). In addition, a short anatomical scan (TR 15.9 ms, TE 4.6 ms, flip angle 27 degrees, FOV 256 x 160 x 216 mm, reconstructed matrix size 256 x 256, isometric 2 mm voxels) was acquired at the beginning of the second and third experiments to facilitate the co-registration of the functional images with the anatomy. The total duration of the MRI study was ~60 minutes. All stimuli were presented using the Presentation^®^ software package (Neurobehavioral Systems, Inc., Albany, U.S.A.).

### Target/Hesitation experiment

The main experiment was composed of three conditions: Hesitation, Target and Actor. The Hesitation and Target conditions were the conditions of interest and used the same movies with different instructions. The movies show an actor grasping one of two balls that are hidden from the viewer inside a box (similar to ref (Thioux and Keysers 2015)). Twelve actors (6 females and 6 males) were videotaped sitting behind a table with only the torso and the hands visible on the video (examples on Fig. 2). Two balls of different size and colour were placed inside the box and the actors were requested to grasp one of the two balls by either telling them to grasp the small or large ball (simple instruction), or by telling them to grasp the ball with the less or more saturated color despite the balls having similar saturation (difficult instruction). In some trials, the actor thus confidently grasped one of the balls, while in others they often hesitated. For each actor six short movies were selected so as to form a block of 6 grasping actions on which to perform the Target or Hesitation task. The individual videos were always cut 3 to 5 frames (120 to 200 ms) before the ball was grasped, and the outcome of the grasping action was therefore hidden from view. Movies were selected to creates blocks that varied in terms of how often the big ball was grasped and how often the actor hesitated.

To ensure that the task requires a careful inspection of the action kinematics, we had to ensure that movie length or the side towards which an actor grasps couldn’t be used to solve the task. The side in which participants grasped was randomized across conditions, with the target on the right side at least once and maximum five times out of six. Hesitation movies were selected by the experimenter on the basis of the visibility of the hesitation and the length of the movies. Hesitations were characterized by a change in reaching direction, or by a pause or slowdown during or at the beginning of the reaching movement. Actions containing hesitation had therefore a specific kinematic profile deviating from the smooth actions without hesitation, however, they also tended to take longer. In order to avoid that the participants could take a decision on the basis of the length of the hesitation movies alone, we selected movies without hesitations that had the same duration than those with hesitation, so that in the final stimulus set, the duration of movies was no longer diagnostic of the number of hesitations, and movie duration could not be used to solve the hesitation task. The Target task also could not be solved based on movie length, because we compared the duration of movies in which the big or small ball was the target, and found no significant difference (p=0.1402, t(136.87) =1.4837). The average length of a video representing a single action was 1440 ms (average 36 frames, Min 17 and Max 72 frames, with a sampling rate of 25 frames/s). Blocks for the fMRI experiment were constructed by pasting the six selected videos together for each actor. The duration of the cross between the videos was set to reach a constant length of 12s for every block.

Participants were requested to answer a question about every block. In the Target condition, they were asked to estimate how many times (out of six) the actor was to pick the big ball. In the Hesitation condition, they were asked how many times the actor was unsure about which ball to pick. The task was presented to 12 pilot subjects in order to verify that hesitations and ball size could be detected above chance level.

As a control condition, we wanted a similar task but without kinematic information to be able to identify the network triggered by the hand-action kinematics. To control for domain general executive components of attention involved in counting, but isolate kinematic information, in the third (Actor) condition we selected single frames of the 12 actors sitting still behind the box, with their hands on the table. The 72 still frames were randomly mixed and grouped by six with a certain amount of repetition of the same actor (2/6, 3/6, 4/6, or 5/6). The task of the participant was to decide how many times the same actor appeared in a sequence. The duration of presentation of the still frames was paired-matched with the length of the movie clips in the other conditions.

Responses were given on a button-box by selecting one of two numbers presented on the screen at the end of the block. The target number appeared on the right side of the screen in half of the cases, and was the largest number in approximately half of the cases. A short question (“Big Ball?”, “Difficult Task?”, “Same actor?”) was displayed for 2.5 s before every block. The question was preceded by a warning sign presented for 0.5 s. The response period was predetermined to 4 s, and was followed by a rest period of 10 s. The total duration of one block was 29 s, including 12 s of action observation. The 12 blocks times 3 condition were pseudo-randomly split over 3 runs, ensuring that each run contained exactly 4 blocks of each condition, that the same actor never appeared in two blocks of the same run, and that the same condition never appeared more than twice in a row. Each run lasted for 6 minutes during which 185 volumes were acquired. Stimuli were back-projected on a tinted screen positioned at the end of the tunnel, and sustained a visual angle of about 15 degrees. The subject’s responses were recorded by the computer used to display the stimuli. The computer also recorded the occurrence of the pulses sent by the scanner at the beginning of each new volume acquisition.

### Action execution experiment

For the action execution experiment, participants were requested to grasp a ball on the left or on the right side of their body upon hearing a low (340 Hz) or a high tone (440 Hz) presented binaurally for 500 ms through magnetic compatible headphones. The two balls (10 cm diameter) were placed at the level of the hips. The participants were requested to reach and grasp the ball, hold it for one second, put it back, and come back in a resting position with the hands on their stomach. The inter-stimuli interval was fixed to 18 s, and the experimenter manually pressed a key on the computer as soon as the subject was back in the resting position to determine the actual length of the execution period. There were 15 grasping actions on each side. The side of grasp was randomized, with no more than 3 consecutive grasps on the same side, and about half of the participants grasping first on the right side. The duration of the execution session was 9 min 30 s during which 285 volumes were acquired.

### False-beliefs / photographs experiment

The false-beliefs/false-photographs task designed by Rebecca Saxe (Saxe et al., 2003) provided an independent localizer for the mentalizing network in the same subjects. In this task, participants read stories followed by a brief statement and have to decide whether the statement is true or false. Half of the story-statement pairs concerned the false belief of a story protagonist, and the other half was about a false representation of the current reality (e.g. on an old map or photograph). For each participant, 12 false-beliefs and 12 false-photographs stories were randomly selected from an initial set of 48 stories, translated into Dutch by native speakers. The 24 stories were divided in two sets with 6 of each story type per block and an equal number of true and false statements per category. The two sets of stories were presented in two successive runs, in randomized order with no more than 3 stories of the same type in a row, and no more than 4 identical responses (true or false). A warning sign was presented before each trial for 1 s. The stories were presented for 12 seconds, immediately followed by the corresponding statement for 5 seconds. After the statement, the words true and false were displayed on the screen, and the participants had to press one of two buttons to indicate whether the statement was correct or not. There was no time limit to answer. The average duration for one run was about 6 minutes.

### Data pre-processing

Data were analysed with SPM8 (Wellcome Department of Cognitive Neurology, London, UK; http://www.fil.ion.ucl.ac.uk/spm) implemented in MatLab 7.13 (Mathworks Inc., Sherborn, MA). Functional images were realigned to the first volume of the first session and co-registered to the anatomical scan acquired just before the experiment using SPM8 default parameters. Then, the anatomy was segmented, and functional images were normalized to the MNI standard space (with a final voxel size of 3×3×3 mm and a 5th-degree B-spline interpolation) using the transformation parameters obtained during segmentation. Finally, data were smoothed with an 8×8×8 mm FWHM Gaussian filter.

### Statistical fMRI Analyses

#### Action observation experiment General Linear Model

At the 1st-level, the 3 conditions (Hesitation, Target, Actor), the instruction, and the response period were modelled separately in a factorial design with three sessions and the motion parameters as additional factors. Averaged parameter estimates for each condition were entered in a factorial design at the second level (random effect analysis, one single factor with 3 levels, non-independent measurements with unequal variances, and masked with a binary image from a smoothed average of the 23 grey matter volumes). The participants’ handedness scores and numbers of error per condition were included as cofactors.

#### Statistical Thresholding

Activity related to the three conditions was assessed at the voxel level using t-tests with voxelwise FDR corrections for multiple comparisons (q < 0.05). Voxelwise FDR was chosen over cluster-size correction because the latter has been shown to inflate false positives (Eklund et al. 2016). However, voxel-wise fdr correction leads to t-thresholds that vary from analysis to analysis. To impose similar standards on all comparisons while also controlling for false-positives, we therefore determined the critical t-value for q<0.05 and the critical t-value for p<0.001. In all our cases, the value corresponding to q<0.05 had a t-value closer to zero (i.e. more permissive) than that for p<0.001. We thus use the more restrictive p<0.001 threshold in all cases, but can be confident that this threshold effectively controls the false-positive rate.

#### Action observation experiment Connectivity Analysis

To identify the path through which the mentalizing and pMNS network exchange information, we used the functional connectivity analysis that comparative methodological studies find most robust: the inverse covariance matrix method (Marrelec et al. 2006; Smith et al. 2011). We extracted the time course of activation from key nodes of the pMNS and the mentalizing network in our data, and explored the significance of the partial correlation between these regions after removing variance shared with the other ROIs and nuisance variables. Specifically, here we performed this analysis only for the right hemisphere, because this is the hemisphere most robustly associated with mentalizing in the literature (Schurz et al. 2014) and the only hemisphere in which all mentalizing nodes (including the rTPJ) are represented in our data, and extracted activity from the core regions of the pMNS and mentalizing network. This was done, by identifying local maxima close to the peaks of the group contrast in the contrast Target-Actor for the pMNS nodes and Hes-Target for the mentalizing network (see Table S2 for MNI coordinates). Specifically, for the pMNS we included the two regions where mirror neurons have been recorded in the monkey (area PF and ventral BA6/44) and its visual input nodes (pSTS and V5). For the mentalizing network this included the most classic nodes (rTPJ and mPFC) as well as the IFG that is found in meta-analyses (Mar 2011; Schurz et al. 2014) and was clearly activated in our paradigm. Fourteen participants showed significant activity in all 7 ROIs (at least 30 voxels at p<0.05 uncorrected) and were included in the analysis. As nuisance variables, we included a total of 12 variables. These included the 6 motion parameters, the average white matter and CSF signal to capture noise. These also included four task variables to capture unspecific activity. They were constructed as for the GLM by convolving block predictors with the hemodynamic response function, and included (i) the timing of the instruction, (ii) the response, (iii) the actor blocks and (iv) a final predictor capturing all the movies, irrespectively of whether the task was Target or Hesitation. This was done so that task specific activation that differentiates Target and Hesitation is not regressed out of the sequences and thus provides the main residual signal that will be tracked across the ROIs (see ref (Cui et al. 2014) for a similar approach). The ICOV was then calculated as follows. Assuming that the matrix of plausible connections between the ROIs is sparse, the inverse covariance method (“glasso” implementation in the R Statistical package) leverages the fact that a full set of partial correlations can be computed using the inverse of the covariance (ICOV) matrix (Friedman et al. 2008; Cui et al. 2014). Briefly, each variable ‘i’ (ROIs in this context) is represented as a general linear model (GLM) comprising of all other variables ‘j’, under the constraint that the sum of the absolute coefficients (C_ij_) of the individual regressors be less than a given constant tuning parameter P. If C_ij_ and C_ji_ is zero, then the ij entry of the inverse covariance matrix is zero (Meinshausen and Bühlmann 2006). This Lasso shrinkage method (Friedman et al. 2008) sets many of the entries in the partial correlation matrix to zero as a function of P. Note that if P is very small the constraint has no effect and a full inverse covariance matrix (and hence a full partial correlation matrix) is obtained. But a small positive P sets many of the partial correlation values to zero, while resulting in different fitting errors for the model. We present the results corresponding to P = 0.025 but the results remain robust against slight variations of this value and are essentially unchanged at P=0.01. The tuning parameter has the effect of controlling the number of predictors in the GLM. The ICOV was calculated separately for each participant, by concatenating the demeaned data from all 3 runs. We then performed a t-test comparing the resulting values for each combination of ROIs against zero, and only those that remain significant after fdr correction for multiple comparison at q<0.05 are considered significant, and are included as potential path in the final connectivity graph.

#### Action execution localizer experiment

For the execution task, the tone and the execution period were modeled separately at the subject level, taking into account the side of grasp. The actual duration of the grasping action was used to determine the number of volumes to be included in every execution block and in the following rest period. One subject was excluded because responses were not recorded. Three other subjects were excluded because of excessive motion. For the rest of the subjects, realignment parameters were included as cofactor and volumes with framewise displacement (FD) above 1.5 mm were neutralized using one regressor per contaminated volume. The parameter estimates for left grasp and right grasp were entered in a factorial design at the second level (non-independent measurements) with handedness scores as a covariate. The execution map showing the areas active above baseline during either left-or right-hand grasp (q < 0.005 FDR). We used a more conservative q value here compared to the rest of the manuscript because action execution generates very strong activations.

##### Mentalizing network localizer experiment

For the false-beliefs/false-photographs experiment, at the 1^st^-level, the story and the statement periods were modeled together (duration 15 seconds) for the two conditions. The warning sign and the response periods were modeled with two other factors and the realignment parameters included in the model (adding one regressor to neutralize volumes with FD > 1.5 mm). One participant was excluded for excessive motion (mean FD > 0.5 mm and more than 5 contaminated volumes). Three other participants did not perform the experiment, two because they were not native Dutch speakers, and one because of a technical problem. The parameter estimates for the two conditions (false-beliefs and false-photographs) were brought to the second level and voxels specific to false-beliefs identified using a t-test comparing the two conditions at FDR q<0.05.

## METHODS STUDY-2: rTMS

### Participants

We screened an initial 30 participants for their resting motor threshold and suitability for MRI and TMS. Because previous experiments had revealed that our TMS system tended to overheat if rTMS was applied with intensities above 65% of the machine’s maximum capacity, and we need to apply TMS at 90% of the passive motor threshold, we selected participants with passive motor thresholds below 72% (i.e. so that 90% of that passive motor threshold would remain below the 65% point at which the rTMS application would lead to overheating). Eighteen participants were selected based on not having counter-indication for experiments involving MRI and TMS, and having a passive motor thresholds ≤72% of our machine output ((mean motor threshold: 57% ± 6%, range: 39% - 65%) and were therefore included in this study. Of these 18, one participant dropped out of the experiment after the first rTMS session, two participants were excluded due to excessive head motion during the rTMS stimulation (as monitored by the neuro-navigation system) and one participant was not able to perform the behavioral task above chance level. The 14 remaining participants (8 females, 6 males) included in the statistical analyses had a mean age of 22 ± 2 years (range: 19-27 years) and a mean handedness of 70 ± 35 on the Edinburgh Handedness Inventory. None of the participants had taken part in Study 1. The experiment was a within-subject design divided in four sessions completed on four different days. During the first day, participants underwent anatomical and functional MRI scans to localize the mentalizing and putative mirror neuron system and identify the target locations for subsequent rTMS. On the same day, the passive motor threshold was determined and participants were trained on the behavioral task. During the last 3 sessions, 15 minutes offline repetitive TMS stimulation was applied to one of the three sites (right PF, right TPJ, vertex) immediately followed by the behavioral task, which lasted for about 6 minutes and was therefore performed while under the effects of the stimulation.

### MRI scan and functional localizer

Functional MRI localizers were used to determine the stimulation sites for each participant individually. Participants underwent one anatomical and four functional runs. First, a high-resolution, structural 3D spoiled gradient image (170 slices; scan resolution = 256 × 256; FOV = 232 mm) was acquired for image co-registration. The four functional imaging runs were conducted using an echo planar T2*-weighted gradient sequence covering 41 sequential axial slices (echo time = 28 ms; slice thickness = 3.5 mm; flip angle = 70°; repetition time = 2000 ms; scan resolution = 64 × 62; FOV = 224 mm). The first two functional runs consisted of two runs of the false-beliefs / false-photographs task, using the same stimuli and procedures as in the first study to identify the rTPJ. Run 3 consisted of an action observation and run 4 of an action execution task used in previous studies to localize the pMNS (Gazzola and Keysers 2009; Arnstein et al. 2011). Masking the action observation activation with the action execution activation is known to generate peaks of activation in the inferior parietal cortex in all participants (Gazzola and Keysers 2009) and was thus used to independently identify participant’s right inferior parietal pMNS target location. Data analysis was performed in BrainVoyager QX 2.2 (Brain Innovation B.V., Maastricht, The Netherlands), which was the software also used for neuronavigation. For every subject, a surface reconstruction of the cortex at the grey-white matter boundary was used to guide TMS stimulation and monitor the distance between the TMS beam and target point. After iso-voxel transformation, T1 images were manually pre-segmented, underwent inhomogeneity correction and were transferred into AC-PC space. The functional images were slice-time corrected using a cubic spline interpolation. Motion correction, spatial smoothing (8 mm FWHM Gaussian filter) and a temporal high-pass filter were applied using standard settings. The functional images were coregistered to the structural image. A boxcar function convolved with the hemodynamic response function was used to model the experimental conditions and the contrasts of interest were computed with a GLM. **rTPJ:** the statistical map generated from the false belief - false photograph contrast considering the two ToM runs were projected onto the reconstructed right hemispheric surface. For each participant, the right TPJ was easily identifiable as a large cluster spanning over the angular gyrus in the posterior IPL and caudal parts of the superior temporal sulcus. Activation was first thresholded at q*FDR*<0.05, then adjusted until a focal area of activation remained. A so-called target point was set onto this focal activation and targeted during rTMS stimulation. The average targeted Talairach coordinates (mean±sd x=51±6 y=-63±4 z=22±6) are in accordance to the coordinates described using the same set of stimuli in other studies (Saxe and Wexler 2005). **rPF**: activation during the action execution run was thresholded at q*FDR*<0.05 and converted into a binary mask. Then, the action observation run was visualized using the complex action – complex control contrast (Arnstein et al. 2011) and masked inclusively with the binary mask generated from the execution run. We then selected the peak of activation within the supramarginal gyrus, corresponding to area PF/PFG where mirror neurons are found in monkeys(Rozzi et al. 2008), as a target point. The average targeted coordinates (mean±sd x=56±5 y=-32±4 z=40±6) fall into the PF complex in regions consistently found to be involved in hand observation and imitation (Keysers and Gazzola 2009; Caspers et al. 2010; Molenberghs et al. 2012). The **vertex** was included as a control stimulation site and its target point was set to the most medial portion of the right postcentral gyrus(Jung et al. 2016).

### Passive motor threshold and training

The passive motor threshold was measured to determine the intensity of the TMS stimulus required for each participant. An electromyogramm was measured from the right first dorsal interosseous (MP36 BIOPAC Systems Inc., Goleta, USA; 43 × 35 mm Hydrogel electrodes). We used a Magstim Rapids^2^ timulator and a figure of eight 70mm coil (The Magstim Company Ltd, Whitland, UK). The starting intensity was set to 50% machine output and single TMS pulses were applied to the approximate position of the left primary motor cortex. The output intensity was increased in 5% steps until a motor evoked potential (MEP) was recorded. The coil position was adjusted to the position of maximal motor evoked potential amplitude. At this optimal scalp position, the intensity was fine-tuned until it met the passive motor threshold criterion (50% of MEPs reaching at least a 50 μV peak-to-peak amplitude). The maximal acceptable motor threshold level was set to 67% machine output (see section: Participants). The motor evoked potentials were recorded in AcqKnowledge 4.1 (Biopac Systems, Inc., USA) using a 30 – 250 Hz Notch filter and a sampling rate of 5000 kHz.

Afterwards, a short training of the behavioral task (described in detail below) was provided. The aim was to get the subject familiar with the design and operation of the task, so that the performance during the first rTMS session would not be influenced by the participant being unfamiliar with the task. We presented one block of five videos of each condition and asked the participant to answer the associated questions. None of these videos appeared in the actual experiment.

### Stimulus creation

A new set of movies with higher definition (50 fps) were shot for this study, involving 10 actors performing the same grasping task under the easy (ball size) and difficult (saturation) conditions as in the first study. Ninety videos (duration 18-116 frames) were selected from the raw material, 9 videos per actor. The videos were selected on the responses of 3 raters familiar with the task who judged whether the actor was given the easy or difficult instruction. We only selected videos with 33% or better accuracy across the raters. On every movie clip, a letter (q or x) was then superimposed on the torso of the actor for 1 to 3 frames in the middle frame of that movie (corresponding roughly to the time when the hesitation and hand aperture also become evident). In the control condition, participants had to detect the presence of the letter q in the movie clip (Figure 2).

From the 9 movie clips per actor, three were attributed to each of the three tasks so as to achieve similar target performance levels of 70%-75% for each task to be above chance level while avoiding ceiling effect. A pre-test conducted with 6 subjects unfamiliar with the experiment yielded performances (mean ± SD) of: 70% ± 9% (Target), 74% ± 6% (Hesitation), 72% ± 10% (Control). This level of performance in the Target task is due to the fact that the ball that is being grasped is hidden in the box, and participants need to anticipate the size of the ball from subtle differences in hand aperture. Similarly, in the other tasks, we achieved this performance by making the letter small and short, and the Hesitations subtle. Examining the duration of the single movies used in this task revealed that there was no difference in duration between Big ball and Small ball movies (p=0.33) or between q and x letters (p=0.65), but the movies with hesitation (average 2.5s) where longer than those without hesitation (average 1.2s, p=0.0002).

The same set of videos was used for all three rTMS sessions, because stimuli like natural movements cannot be used to create three perfectly matched sets of videos for the three rTMS target sites. Although, the appearance order of videos was randomized for each session, a significant reduction in response latencies over sessions was observed.

### rTMS sessions

This study was set-up as a repeated-measures design. Each participant performed three tasks (Target, Hesitation, Control) following rTMS stimulation on three different stimulation sites in the right hemisphere (TPJ, PF, vertex) on three different days. The order of stimulation sites was counterbalanced across participants. The minimum time between two rTMS sessions was two days to avoid the influence of stimulation after-effects. The participant was seated in front of a monitor that was used to display the neuronavigation tracking throughout the rTMS stimulation and the behavioral task afterwards. The TMS coil positioning system (Zebris Medical GmbH, Isny im Allgäu, Germany) was used for neuronavigation by coregistration of the head of the participant to the TMS coil. The TMS coil was fixed in place using a holder and its position relative to the target point monitored using BrainVoyager and the neuronavigation system. The experimenter readjusted the coil position whenever necessary. External cooling was applied to the TMS coil throughout the stimulation. The handle of the coil was pointing backwards at a 45 degree downwards angle for the TPJ and PF sessions and pointing backwards during the vertex session. During the rTMS sessions, the participants received a 900 pulses 1 Hz rTMS stimulation at 90% motor threshold on one of the stimulation sites. Repetitive, low-frequency (1 Hz) TMS stimulation has an inhibitory effect on neuronal activity (Chen et al. 1997; Pascual-Leone et al. 1998; Maeda et al. 2000).

After the stimulation, participants performed the behavioral task, which lasted on average 6.52 ± 0.31 minutes with a range of 5.74 to 7.31 minutes. Thus, the behavioral tasks were all completed within the 7.5 minute time window of inhibitory effect typically considered to be induced by the TMS stimulation we chose(Knecht et al. 2003). One experimental run consisted of a written explanation of the task, the display of 90 videos (30 videos per condition) and the participants answer selection. Importantly, each video included all visual features to complete any of the three tasks (Figure 1). The current condition was indicated before each video by displaying one of the three experimental questions: Big ball? – Hesitation? – Letter q? Videos of the same condition were presented in blocks of five. No more than two blocks of the same condition were presented sequentially. After each video the participant selected an answer in a ‘Yes’ or ‘No’ fashion, according to the question asked (Big ball? – Hesitation? – Letter q?). The question screen was presented immediately after each video, there was no time limit given to respond. Participants used a foot pedal to give their answers to avoid hand movements that could have interfered with hand action observation. To avoid motor preparation during the video, whether the right or the left pedal indicated a ‘yes’ was randomly chosen after each movie, and indicated on the screen. The behavioral task was displayed using the Presentation^®^ 16.1 software (Neurobehavioral Systems, Inc., Berkeley, USA).

### Data analysis

The performance during the behavioral experiment was assessed in terms of task accuracy and response latencies. All statistical analyses were performed using IBM SPSS Statistics 23 (IBM Corporation, Armonk, USA). The threshold for all statistical tests is p = 0.05, unless stated otherwise. Due to the within subject experimental design we expected an effect of experimental session (day1 – day2 – day3), independently of which stimulation site was targeted, because the repetitive usage of identical stimuli was likely to cause training effects. As expected we found a significant decrease in reaction time across days irrespective of task (see results), but found no such effect on accuracy. By randomizing the order of stimulation site, this effect is orthogonal from the effect of stimulation site, but it still inflates residual errors. To remove this training effect, we subtracted the response latencies of the control task from those of the two experimental tasks. One possibility would have been to perform a simple subtraction like in the following formula for response latencies to the Target task after rTPJ stimulation: *RT*(*Goal_TPJ-detrend_*)_*i*_ = *RT*(*Goal_TPJ_*)_*i*_ – *RT*(*Control_TPJ_*)_*i*_ However, this would remove the true effect of rTMS on the Target task if rTPJ sitmulation also affected the control task. To prevent this, we added the average RT difference across the stimulation site and the vertex back onto the corrected values using the following formula for each participant i, for each task and stimulation site:

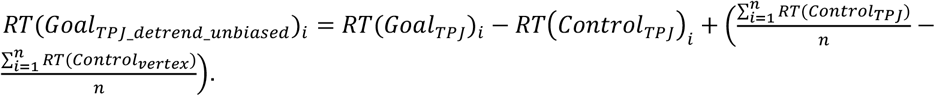

To ascertain whether this procedure reduces noise variance due to inter-individual and between session variability, we calculated the change in variance between the detrended and raw data. The standard deviation dropped by 59% for the Hesitation*TPJ* latencies and by 48% for the Hesitation_*PF*_latencies. The standard deviation dropped by 52% for the Target_*TPJ*_latencies and by 52% for the Target_*PF*_latencies.

## RESULTS

### Study 1 – Activation as a function of condition

Contrasting the Target task against our Actor task in our 23 participants revealed a bilateral network that closely corresponds to the typical pMNS or action observation network reported in the literature (dPM, vPM, PF, BA2, posterior mid-temporal gyrus including human V5, Figure 3a). Contrasting the Hesitation task against our Actor task (Figure 3b) revealed a somewhat similar network. To compare these activations with those usually associated with the pMNS and mentalizing network we used two approaches: literature meta-analyses and localizers in our own participants. Activity is typically associated with the pMNS network if it is triggered by the observation of actions and falls within voxels also involved in executing similar actions (Gazzola and Keysers 2009; Caspers et al. 2010; Rizzolatti and Sinigaglia 2010). As the activity is already triggered by action observation in our contrasts, the critical question is whether the voxels overlap with those involved in grasping. We thus used neurosynth with the search term grasping, and used the *reverse* inference map (i.e. locations in which activations increase the likelihood that participants had been asked to grasp, i.e. p(Grasping|Activation) (Yarkoni et al. 2011). Because many grasping studies only use the dominant right hand, this network was strongly lateralized to the left hemisphere, and lacked the right primary motor cortex. To avoid this bias, we left-right flipped this map, and then used the maximum value in each voxel across the flipped and unflipped map. The resulting map is shown in red in Figure 3c. For mentalizing, we used the meta-analysis by Mar (Mar 2011), which is shown in blue in Figure 3c. Second, we also performed localizers for grasping and mentalizing in our own participants, which lead to conceptually very similar results (Figure S1 in supplementary materials). In Figure 3a,b, we show the activity triggered by the contrast in yellow, and the overlap with the grasping network in red, and that with the mentalizing network in blue. This revealed that the pMNS (red) is recruited in both contrast, while many nodes of the mentalizing network (mPFC, aIFG, STS) are only recruited in the Hesitation task. Comparing the two contrasts directly reveals regions that are specifically (more) recruited by one of the experimental conditions (Figure 3d). The Target task (hot colors in Figure 3d) activates BA2, PF, dPM and vPM (BA6/44) more than the Hesitation task (Table 2). The Hesitation task activated a network (blue in Figure 3d) including mPFC, rTPJ, aIFG, the temporal pole and the superior temporal sulcus (STS) more than the Target condition. The activity that was larger during the Target task (hot colours Figure 3d) fell within the grasping network (hot colours in Figure 3e) but not the mentalizing network (no hot colors at all in Figure 3f), while the activity larger during the Hesitation task fell exclusively (when using the meta-analytical networks) or mainly (when using our own localizers) in the Mentalizing network (cold colours Figure 3f). Because the connectivity analysis was restricted to the 14 participants that had significant activation individually in each of the 7 ROIs explored, we repeated the GLM also for these 14 participants only. This analysis lead to conceptually identical results as for the entire sample (Supplementary Figure S2).

**Figure 3:**
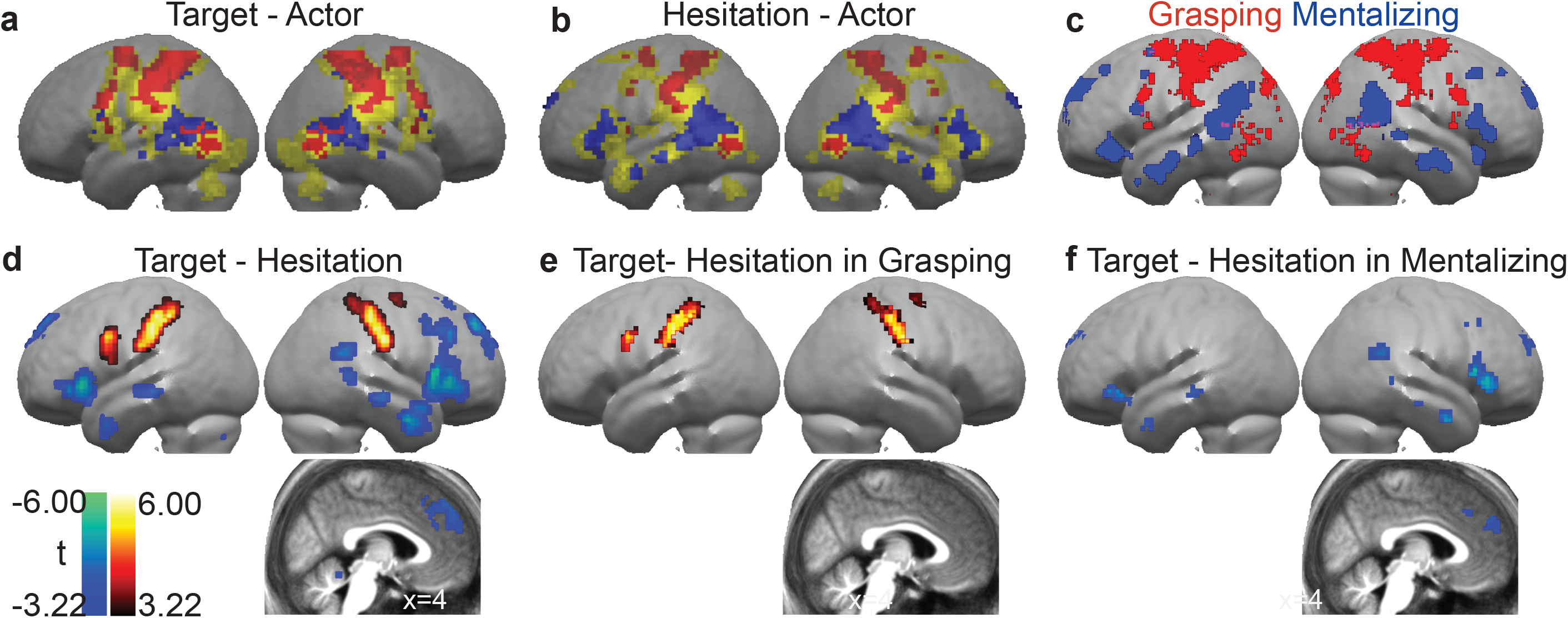
FMRI results. **(a-b)** Results of the basic contrasts against the control condition. Voxels with significant activation are shown in yellow. Those falling within voxels associated with grasping are shown in red, those falling within voxels associated with mentalizing in blue. This color-coding is based on a meta-analysis of papers examining the execution of grasping (red) and mentalizing (blue) shown in panel **(c)**. **(d)** Contrast Target-Hesitation, with warm colours indicating activity larger for Target, and cold colours activity larger for Hesitation. **(e and f)** same as **d** but inclusively masked with the red and blue grasping network of **c**, respectively. All results are shown on renders of the mean normalized gray-matter segment of the participants, and thresholded at p<0.001 uncorrected (t>3.22 or t<-3.22). All results survive q<0.05 voxel-wise false discovery correction for multiple comparison (i.e. the critical t-value for q<0.05 was closer to zero than 3.22, and results shown at t>3.22 thus contain less than 5% false positives, see methods section ‘Statistical Thresholding’. The sagittal slices in d, e and f are taken at x=4 to show the mPFC activation or absence thereof. See Supplementary Materials for a similar analysis based on our own localizers.

**Table 2:**
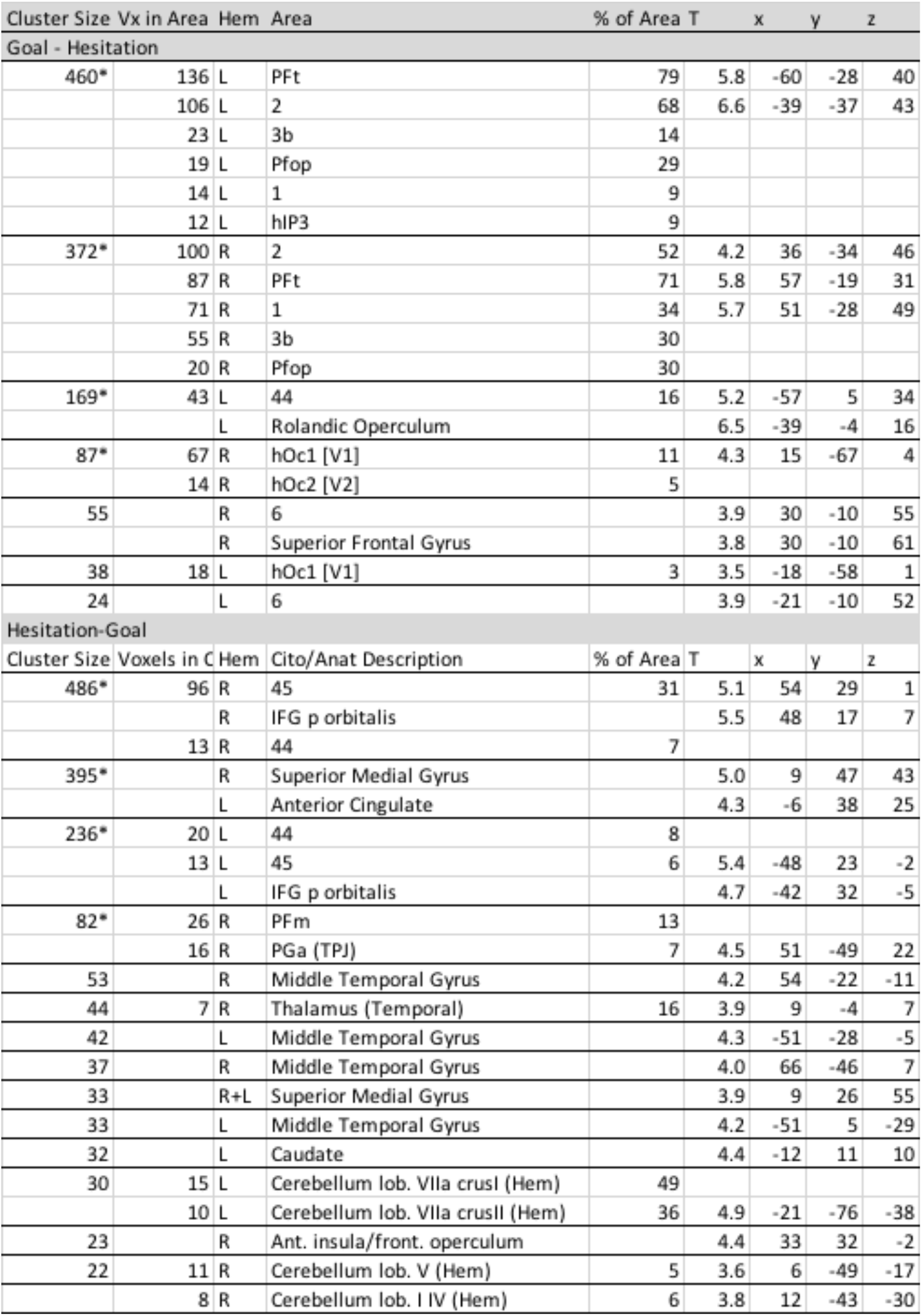
Results of the direct comparison of Target and Hesitation labelled using SPM Anatomy Toolbox. As for Figure 3, only voxels surviving voxel-wise false discovery correction q_fdr_<0.05 are presented (see methods section ‘Statistical Thresholding’ for details), and voxels with t<3.22 (p_unc_<0.001) and clusters of less than 10voxels are not shown. From left to right: the cluster size in number of voxels, the number of voxels falling in a cyto-architectonic area, the hemisphere (L=left; R=right), the name of the cyto-architectonic area when available or the anatomical description, the percentage of the area that is activated by the cluster, the t values of the peak associated with the cluster followed by the MNI coordinates in mm. Note that for regions for which there are no probabilistic maps, an anatomical description is provided, but columns 2 and 5 then have no values. * marks clusters large enough to survive a cluster-level family wise error correction at p<0.05.

### Study 1 - Connectivity Analysis

To explore how information is transferred across these networks, we used the functional connectivity methodology that has been considered most robust in comparative studies (Smith et al. 2011), the ICOV method (Marrelec et al. 2006; Cui et al. 2014). This method calculates the partial correlation across every two nodes of interest in the brain after conditioning out the variance shared with any other nodes and the nuisance variables (see method). As in mediation analysis, the use of *partial* correlations ensures that if A is connected with B and B with C, but A is not connected with C, then partial correlations should reveal significant partial correlation between A-B and B-C but not A-C, whilst traditional correlation would also detect significant correlation along A-C. We restrict our analysis to the right hemisphere, because the GLM analysis revealed significant increase of activity in the right but not the left TPJ and the right hemisphere included the shared visual input regions V5 and pSTS, the core nodes of the pMNS (PF and vPM) and the core nodes of the ‘blue’ mentalizing network preferentially activated during the Hesitation task (Figure 3d, rTPJ and mPFC and anterior IFG). The ICOV was then calculated subject by subject, and significance of each connection assessed by testing whether the partial correlation differs from zero on average in the population using a t-test, Boneferoni-corrected for the 21 possible connections. This analysis revealed two networks, one that includes the pMNS (red) and one that includes the mentalizing network (blue), which are connected via a single route from vPM to aIFG (see Figure 4). In this graph, the blue network does not receive direct visual information from V5/pSTS, but this input is mediated by a network overlapping with the grasping localizer and associated with the pMNS. Specifically, the aIFG serves as a *trait d’union* between the visual information and pMNS on the one hand, and the classic TPJ and mPFC nodes of the mentalizing network, on the other.

**Figure 4:**
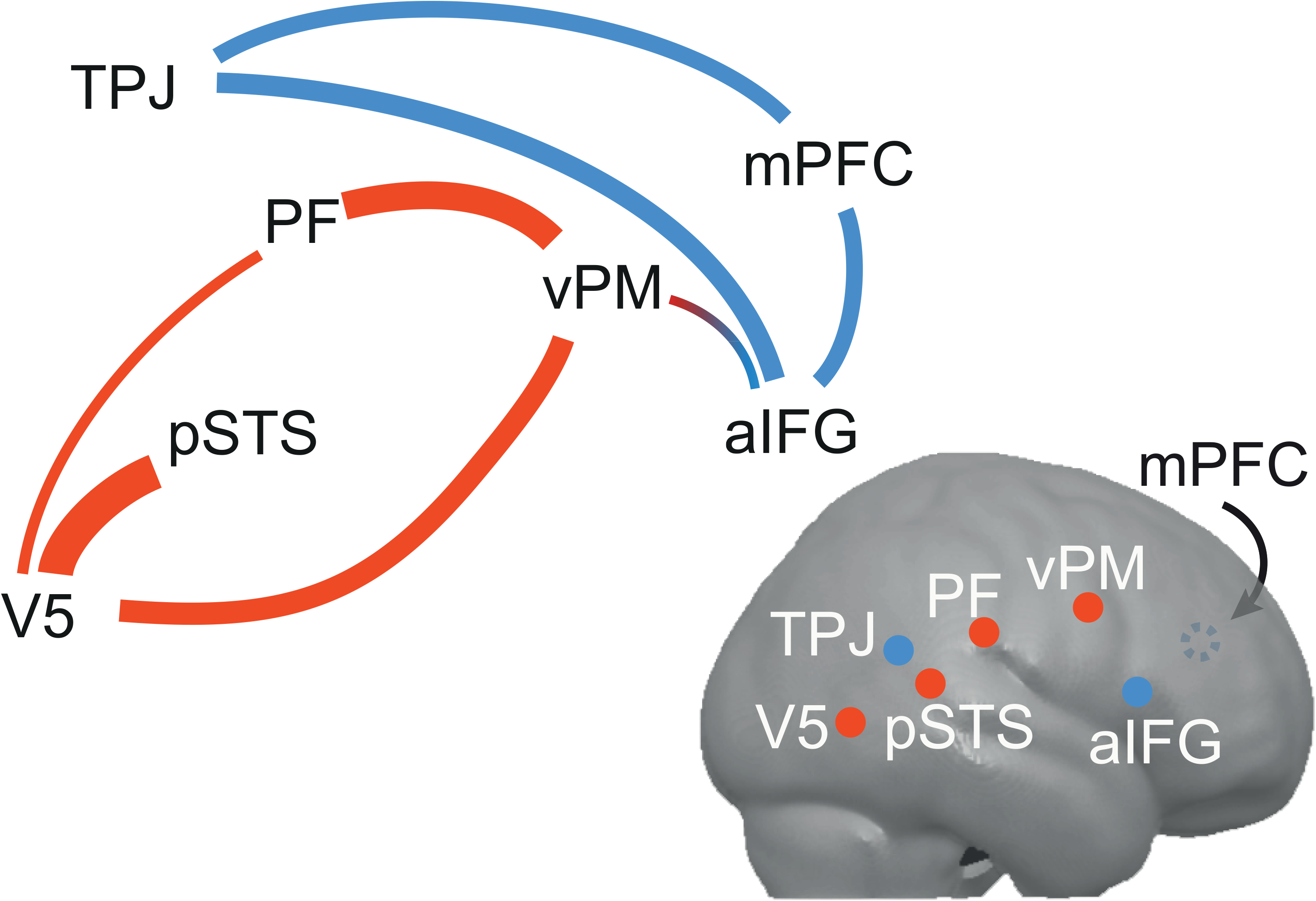
Connectivity graph across the nodes in the right hemisphere of the common network (red) and the mentalizing network (blue). The line thickness reflects the average partial correlation coefficient, and only significant partial correlations are shown (p<0.05 Boneferoni-corrected). The mPFC node is on the medial surface of the brain, and is thus shown as a dotted circle. Note how the anterior IFG node mediates the crosstalk between the two networks.

**Figure 5:**
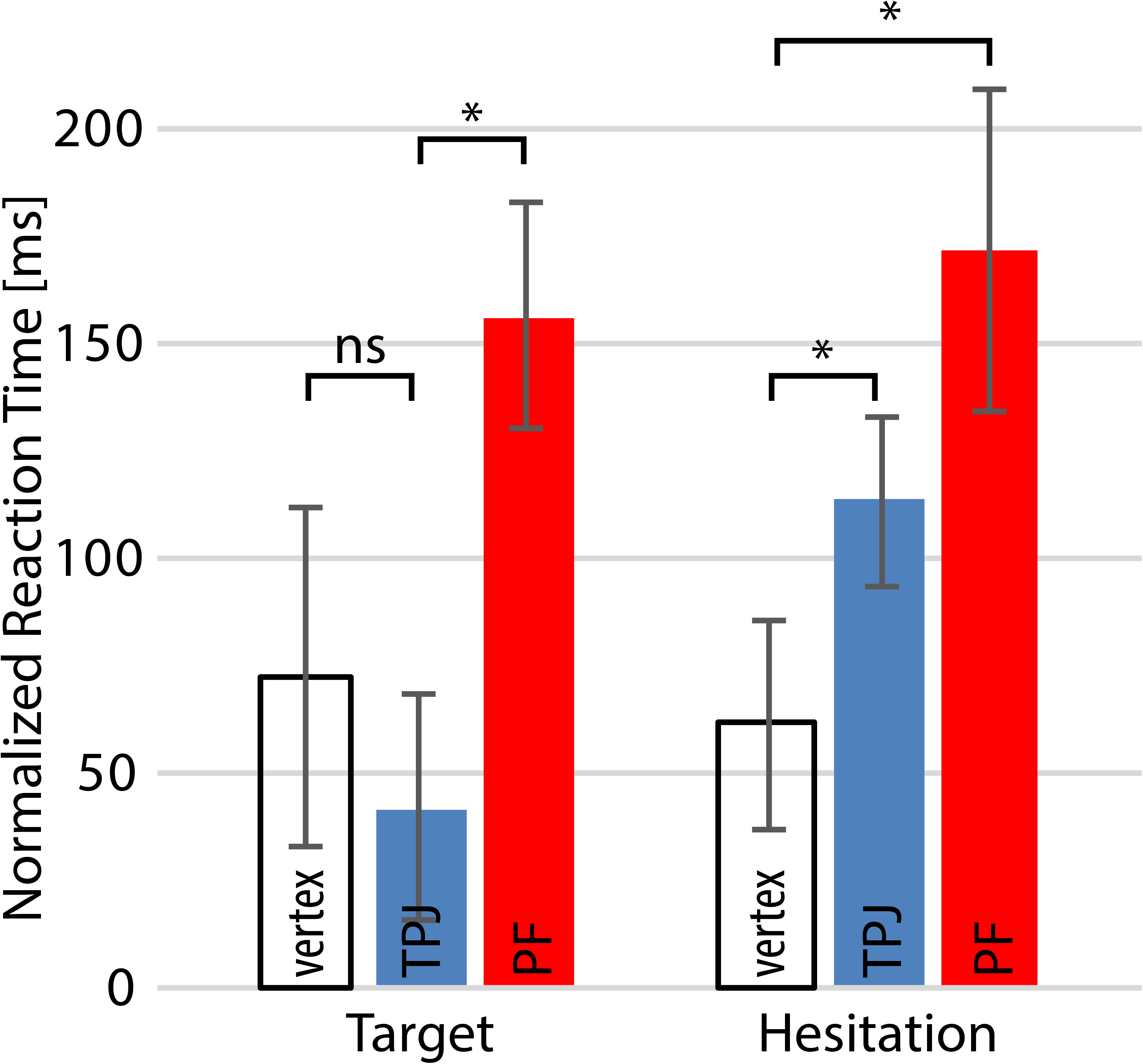
Task-dependent increases in normalized response latencies due to repetitive TMS stimulation of TPJ or PF. The y-axis represents normalized reaction times in ms, from which the reaction times of the control condition have been subtracted (see Methods). There was a significant Task (Target vs. Hesitation) x Stimulation site (vertex, TPJ, PF) interaction (F(2,12) = 4.89, p<0.05), which was further explored using 4 planned comparisons (indicated by brackets). Error bars show the standard error of the mean, * = p < 0.05.

### Study 2 – TMS results

To test the necessity of both the mentalizing and pMNS networks for optimal performance, we conducted a follow-up repetitive TMS experiment. We used an independent localizer task for mentalizing to functionally localize the right TPJ. We selected the right TPJ as one of our target region because this is where we found an effect of Hesitation > Target in our pilot fMRI study (there was no significant increase of activity in left TPJ for Hesitation detection), and because this brain area that can easily be targeted with rTMS (as opposed to MPFC) and has been shown to be consistently recruited by mental state attribution tasks across the literature (Schurz et al. 2014). We used a localizer of pMNS activity to functionally localize the right PF node of this system as our second target region (because targeting vPM leads to discomfort). Finally, we targeted the vertex as a control location. The functional localizers, and the three target locations were performed in a within subject design on 4 different days. On each of the rTMS days, the reaction time was measured while participants determined if the actor in the movies grasped the big ball (Target) or hesitated (Hesitation), and the reaction time of a control task (in which participants had to identify a letter in the movies (Letter) was subtracted from the raw reaction times to remove unspecific inter-individual differences in reaction times, day-to-day variations in fatigue or unspecific motor-output delays caused by TMS application (Figure 4).

To test whether the tasks were matched in terms of difficulty, we employed a 3 stimulation sites (TPJ, PF, vertex) x 3 tasks (Target, Hesitation, Control) repeated-measures ANOVA for task accuracy. There was a non-significant trend for stimulation site (F(2,12) = 3.45, p = 0.0655), no effect of task (F(2,12) = 0.22, p = 0.8) or interaction between stimulation site and task (F(4,10) = 1.05, p = 0.43). The three tasks were thus matched in terms of difficulty (accuracies (mean±SD): Target: 74% ± 6%, Hesitation: 76% ± 8%, Control: 81% ± 10%), and preselecting movies to achieve a performance around 70-80% had been successful. This performance level is often ideal to reveal subtle TMS effects.

To test whether rTMS had an effect on latency, a 3 stimulation sites x 2 tasks within subject ANOVA was performed using the multivariate approach in SPSS on the normalized latencies (i.e. latencies relative to the control task, see methods, and Figure 4). This revealed a significant effect of stimulation site (F(2,12) = 4.20, p=0.041, partial eta squared = 0.41) and a significant interaction effect of stimulation site and task (F(2,12) = 4.89, p=0.028, partial eta squared = 0.45). We had two specific hypotheses to test via planned comparisons: for the Hesitation task, both PF and TPJ stimulation should lead to slower responses than vertex. This was true: PF and TPJ stimulation lead to slower responses than Vertex (t(13) = 2.4, p=0.016; t(13) = 2.3, p=0.02). In addition, for the Target task, we expected the PF stimulation to disrupt performance more than TPJ stimulation, and TPJ stimulation not to have an effect: both were the case (PF vs. TPJ: t(13) = 3.1, p<0.005; TPJ vs. vertex: t(13)=0.9, p=0.19). Only two planned comparisons were performed per task (n-1, where n is the number of factor levels) to avoid inflation of false positives. A similar trend is visible in the raw latencies (Supplementary Figure S3). Excluding the PF condition and performing a 2 (vertex vs TPJ) x 2 (Target vs. Hesitation) ANOVA confirmed a significant interaction (F(1,13) = 8.36, p=0.013, partial eta squared = 0.39), corroborating the fact that the interaction in the 3 x 2 ANOVA was driven by the fact that TPJ stimulation indeed has a significantly higher impact on the Hesitation than Target task.

## DISCUSSION

Using fMRI, we confirm that when participants process very physical mental states such as whether a hand is to grasp a large or small object, they recruit the action observation network traditionally associated with the pMNS including visual regions in the pSTS, V5, parietal areas PF and the primary somatosensory cortex (SI, BA2 in particular), as well as frontal regions in the ventral and dorsal premotor cortex (vPM and dPM). When viewing the same movies to determine a more abstract mental state that remains closely linked to action kinematics, hesitation, they recruited a network associated with mentalizing including the rTPJ, mPFC, and aIFG. This activation of that mentalizing network co-existed with a significant activation of the pMNS. To explore whether the pMNS and mentalizing network worked independently during this task, or whether the mentalizing network built upon the computations of the pMNS in our Hesitation task, we performed a mediation analysis to explore whether the timing of the BOLD signal in the mentalizing network is better explained using direct connections with V5 and pSTS, or via the pMNS. We used a method based on partial correlation because this method had been shown to be particularly robust and to have high sensitivity to detect network connections (Marrelec et al. 2006; Smith et al. 2011). This revealed that information from the core biological motion nodes (V5 and pSTS) is exchanged with the mentalizing nodes (mPFC and TPJ) via the pMNS (PF and vPM), with the anterior IFG serving as a connection between the pMNS and the mentalizing network. This pattern of brain activation and connectivity suggests that disrupting the pMNS should interfere with both the Target and Hesitation task, while disrupting the mentalizing network should interfere only with the Hesitation task. As hypothesized, in a follow-up experiment using TMS we found that performance in the Hesitation task was susceptible to interference with both networks: interfering with TPJ or PF in the right hemisphere lead to slower reaction times than vertex stimulation. On the other hand, performance in the Target task was only susceptible to interference with the pMNS network: interfering with PF lead to slower reaction times than interfering with TPJ, that in turn did not differ from interfering with the vertex.

A small number of studies corroborate our fMRI findings by using photos or movies of actions as stimuli and observed both networks to be co-activated, namely while judging the meaning of gestures (Schippers et al. 2010), the confidence of an informant from facial movements (Kuhlen et al. 2015), and while judging why participants perform certain hand actions (de Lange et al. 2008; Spunt et al. 2011; Ampe et al. 2014). Co-activation were also found, when participants observed actions directed at another person as compared to solo actions (Centelles et al. 2011; Becchio et al. 2012). Our findings shed light on why some studies find co-activation while others do not. By contrasting the Target and Hesitation task on the same stimuli we show that the pMNS is recruited whenever participants view goal directed actions in our sample, but that the mentalizing system is additionally recruited only when participants are asked explicitly to process mental states that go beyond the immediate motor target of the action. By performing connectivity analyses, we add to these studies by showing that while processing mental states that go beyond the target of the action but that can be inferred from the kinematics of the action such as hesitation (Patel et al. 2012), the pMNS seems to mediate (at least in part) the interaction between the mentalizing network and visual information from V5 and pSTS, and that this exchange of information between the pMNS and the core nodes of mentalizing (rTPJ and mPFC) occurs via the anterior IFG. A careful comparison between the cytoarchitectonic map of BA44 with the locus of that IFG cluster indicated that over 90% of it was outside and anterior to BA44 (including BA45 and BA47). This pattern of connectivity is in line with what we know about the anatomy of the connections across these systems in macaques, in which two separate fronto-parietal networks exist, that have homologies to the pMNS and mentalizing network we find here in humans, respectively. The first connects V5 and pSTS via reciprocal connections with parietal area PF/PFG which in turn are connected to the ventral premotor cortex (area F5) (Nelissen et al. 2011). The second connects more anterior and ventral prefrontal regions (homologue to human BA45/47) with the medial prefrontal areas 9 anteriorly and cortices at the temporo-parietal interface including area PG and adjacent regions of the temporal sulcus (Yeterian et al. 2012), posteriorly. Importantly, these two networks are interconnected via direct connections between F5 and BA45/47 (Yeterian et al. 2012). Comparative studies confirm a similar pattern of connectivity in humans (Mars et al. 2012; Thiebaut de Schotten et al. 2012). These anatomical findings therefore fully concur with our functional connectivity results that reveal two separate functional networks, interconnected via the anterior IFG, and provide support for our functional connectivity analysis.

Our finding that rTMS over rPF and rTPJ lead to a slowing of reaction times for the Hesitation task provides what is to our knowledge the first evidence that both the pMNS and mentalizing networks are necessary to optimally attribute the same mental state (hesitation) to an actor from observing his actions. Because rTMS has limited spatial resolution, it is important to ascertain that our rTMS stimulation on rPF had not simply spilled into rTPJ and vice versa, but that rTMS on rTPJ and rPF had different effects in the Target task speaks against this possibility. Given that interfering with activity of a specific node of the pMNS leads to changes across the entire pMNS (Valchev et al. 2016), we advise against concluding that rTPJ and rPF are necessary for optimal performance in our Hesitation task. Instead our data suggests that the pMNS and metalizing networks (of which the rPF and rTPJ are part) are necessary for the optimal attribution of hesitations. Recent meta-analyses have confirmed that disruptions of the premotor and parietal pMNS nodes via neuromodulation (Avenanti et al. 2013) or lesions (Urgesi et al. 2014) leads to impairments in tasks that resemble our Target task, but these studies did not explore or reveal contributions of the mentalizing network to interpreting mental states from such observed actions. Similarly, reviews confirm that a range of studies has implicated the TPJ in mental state attribution (Donaldson et al. 2015), but that region had so far not been systematically investigated in relation to attributing mental states from observed actions.

Our results are corroborated by those of another rTMS study showing that interfering with the pMNS impairs the ability to detect lies on the basis of body kinematics (Tidoni et al. 2013), however that study failed to find an effect of TPJ stimulation. This contrasts with our finding that rTMS to the right PF and TPJ interferes with hesitation attribution. In this earlier study however, authors targeted the left hemisphere, and the stimulated areas were not defined on the basis of the participants fMRI data. Our data reinforces this finding by showing that both networks interact functionally and are both necessary for optimal inference of mental states from observed actions.

### Nature of the computations leading to the detection of hesitation

Our experiment did not systematically study what stimulus properties lead to the detection of hesitations. However, the kinematics of the grasping action when actors reported to be confident and were perceived as confident are clearly smoother and on average faster than the kinematics in the less confident conditions, and previous studies have shown that the kinematics and durations of actions suffice to determine confidence (Patel et al. 2012). Accordingly, a simple way to compute an estimate of hesitation would be to (a) compute a prototypical energy-efficient kinematic for grasping the balls in the box, and (b) compare that prediction with observed kinematics. The larger the deviation, the higher the likelihood that the actor was hesitant. The somatosensory-motor grasping network has been shown to be essential for drawing non-mentalistic inferences from observed kinematics of a grasping action (Pobric and de C. Hamilton 2006; Michael et al. 2014; Avenanti et al. 2017; Valchev et al. 2017), and the predictive coding models suggested to be implemented in the pMNS could provide an architecture to calculate the deviation from such prediction (Grèzes et al. 2004; Keysers and Perrett 2004; Kilner et al. 2007; Keysers and Gazzola 2014; Keysers et al. 2014; Avenanti et al. 2017; Thomas et al. 2018). This could explain why interfering with the pMNS slowed the ability to detect Hesitations in our data, and why this network is activated and feeds into the mentalizing network. An involvement of the motor system in detecting hesitation is also predicted by the elegant observation that confidence ratings from observed actions can be predicted by the viewers own motor execution of a similar task (Patel et al. 2012). Alternatively, the visual system alone could have computed smoothness and velocity of movement and forwarded that information to the mentalizing network. However, if this were sufficient, why would rTMS over the pMNS slow reaction times, and why would the pMNS seems to mediate part of the functional connectivity between visual motion areas and the mentalizing network? Importantly, that interfering with rTPJ slowed reaction times in our Hesitation task more than in our Target task indicates that attributing hesitations to others seems to require something in addition to the visuo-motor computations also involved in extracting the intention-in-action (i.e. what object is the target). Given that activations in the rTPJ support reverse inference for mentalizing (see reverse inference map for mentalizing in neurosynth.org), this suggests that this additional and necessary computation may indeed pertain to the fact that hesitations are mental states more abstract than those processed by the pMNS and may share some operations with those recruiting rTPJ during more typical mentalizing tasks. It remains unclear what these operations involve from a neuro-computational point of view, which commands caution in such reverse inference. However, our conclusion is consistent with the results of a recently published study (Chambon et al. 2017) that found the backward connectivity between MPFC and right TPJ to be increased when processing higher levels of intention (an object is moved to build a shape vs. is simply moved), and to be influenced by the priors of the subjects (how likely one action was to occur).

### Limitations

Our study has limitations that should be considered while interpreting its results. First, there has been a long tradition of associating brain activity in regions such as PF and vPM with the pMNS and simulation due to the homologies with regions containing mirror neurons in monkeys and the fact that this activity falls within voxels also activated during the execution of grasping in our subjects and in meta-analyses. Here we follow that tradition to link to that substantial literature. However, it is important to realize the limitations of such reverse inference, even when, as we did here, a reverse inference meta-analysis is used (i.e. p(Grasping|Activation) rather than p(Activation|Grasping) (Gazzola and Keysers 2009; Poldrack 2011; Yarkoni et al. 2011)). The same limitation applies to interpreting activity, or the effects of a disruption, in TPJ to mentalizing. Accordingly, the labels pMNS and mentalizing networks should be taken as tentative labels for large scale networks rather than strong claims on the function underlying the activity in these regions. This is similar to the use of default mode (Fox et al. 2005) or saliency network (Seeley et al. 2007), that do not suggest that all activity in these networks reflect default processes or saliency computations.

Although we have chosen one of the most robust functional connectivity methods (Smith et al. 2011), our findings that the pMNS mediates the visual information exchange between V5/pSTS and rTPJ and mPFC in the right hemisphere remains restricted to our particular choice of ROIs. Accordingly, our findings do not exclude that rTPJ/mPFC receive some form of direct visual input, but rather than the pMNS appears to provide some of the input to the mentalizing system. Indeed, relaxing the regularization parameter in our ICOV analysis to allow more connections to remain significant leads to the emergence of connections between visual regions and the mentalizing network (data not shown), reinforcing the notion that our result should be interpreted as suggesting a mediation by the pMNS (which is true independently of the regularization parameter) rather than the absence of any direct input (which is true for some but not all parameters). Interestingly, this mediation remains evident even if the task predictors are removed from the ICOV analysis (data not shown). A strength of our study is that the weaknesses of a specific methodology (e.g. fMRI and connectivity analysis) is supplemented by independent evidence with a different methodology (rTMS) that also suggests that the pMNS network contributes to the optimal performance of the Hesitation attribution task.

When applying rTMS over a brain regions involved in hand actions, as we did in this study, it is important to exclude the possibility that a slowing of reaction time simply reflects an unspecific motor output impairment. To mitigate this risk we compared reaction times in the control task (Letter detection) across the three stimulation sites, and found no significant differences (F(2,26)=0.01, p=0.99)) suggesting that rTMS over rPF did not slow down motor output. In addition, we did not compare raw reaction times across stimulation sites, but the differences in RT between the tasks of interest (Target or Hesitation) and the control (Letter) task to remove any residual non-specific effects on motor output.

Finally, our control condition in Study 1 differs from the other conditions in the absence of movement in the stimuli and in the detection of a different property (identity). This must be considered when interpreting the results of contrasts between this control task and the Target and Hesitation task. The shared network between these two tasks (as compared against the control task) could in theory contain regions not activated in both experimental conditions but inhibited in the shared control condition. We therefore performed the same analysis using the implicit baseline instead of the control condition, but this revealed results conceptually identical to those presented in Figure 3. In fact, relative to fixation the control condition with still frames of the actors was also associated with a significant increased activity in the pMNS network (Supplementary Figure S4).

## CONCLUSIONS

Our findings refine our understanding of the relationship between two of the most studied neural networks in social cognition: the putative human mirror neuron system and the mentalizing network. Using a new paradigm in which participants view goal directed hand actions that permit both the identification of immediate intention (which object will be grasped?) and more abstract mental state (does the actor hesitate?), and using a combination of brain measurement and modulation we were able to constrain the contributions of the two systems to action comprehension. When participants see the kinematics of other people’s actions, and extract abstract mental states from those kinematics (as opposed to the immediate intention-in-action), both systems are recruited, information is exchanged across them and most importantly, both are necessary for optimal mental state attribution. Hesitation detection is probably relevant from an evolutionary point of view. Detecting others’ hesitations can be useful to identify the difficulty of a task and the time it might take to obtain a reward, it may give the advantage in a competition, revealing the hidden intentions or weaknesses of an opponent. Hesitation detection does not require language, hesitation is not a propositional attitude, and the detection of hesitation-in-action can be readily achieved by an analysis of the discrepancy between perceived and expected action kinematics. Still, our study shows that this pre-processed motor information feeds into a mentalizing network that is essential for associating deviations from a predicted motor program with a specific mental state. Hesitation detection could be a phylogenetic precursor to the ability to represent others’ mental states.

## Acknowledgements

We are grateful to Valeria Gazzola for the original idea of searching for the missing link between mentalizing and MNS and for helping us to refine the experimental design, to André Aleman for lending us the rTMS apparatus, to Rajat Thomas for the ICOV analysis, to Nicolas Valchev for help with the rTMS setup, to Michel Weber for discussions about hesitation and to Ysbrand van der Werf for advices on the rTMS protocol.

